## Supplementary material for "Modelling transcription with explainable AI uncovers context-specific epigenetic gene regulation at promoters and gene bodies": S1 Appendix

### S1 Appendix. Supplementary methods

#### Interpretable machine learning techniques

Interpretable machine learning aims to make complex predictive models more transparent by providing insights into how they generate predictions. As machine learning models, particularly deep neural networks and ensemble methods, grow in complexity, their decision-making processes often become opaque, making it difficult to understand the contribution of individual features [1]. This lack of interpretability is a critical limitation, especially in fields like computational biology, where understanding feature importance can reveal underlying biological mechanisms. Various interpretability methods, including feature attribution techniques, surrogate models, and rule-based explanations, have been developed to bridge this gap. These approaches enable researchers to analyse model predictions, assess their reliability, and extract meaningful insights. Among these methods, SHAP [2] offers a principled approach to quantifying feature importance based on cooperative game theory [3], ensuring fairness and consistency in feature attributions.

##### Shapley Additive Explanations (SHAP)

Interpreting complex machine learning models often relies on additive feature attribution, where the model's prediction is approximated as a sum of individual feature contributions. Formally, an explanation model $g$ is defined as a linear function :

$$f(\mathbf{x}) = g\left(\mathbf{x}'\right) = \phi_0 + \sum_{i=1}^{M} \phi_i x_i'$$

where $x'$ represents a simplified input space $x = h_x(x')$, $M$ is the number of simplified input features, and $\phi_i$ denotes the contribution of each feature. This framework ensures that the sum of feature attributions approximates the original model's output $f(x)$. SHAP values are inspired by *Shapley values*, originally developed in cooperative game theory to fairly distribute a total payoff among a set of contributors. Given a predictive model $f$ and a feature set $\mathbf{x}$, the SHAP value for feature $x_j$ quantifies its contribution to the prediction $f(\mathbf{x})$ by computing the average marginal effect of adding $x_j$ across all possible feature subsets:

$$\phi_i(f, x) = \sum_{z' \subseteq x'} \frac{|z'|!\,(M - |z'| - 1)!}{M!}\left[f_x\left(z'\right) - f_x\left(z' \backslash i\right)\right]$$

where $|z'|$ is the number of non-zero entries in $z'$, and $z' \subseteq x'$ represents all z' vectors where the non-zero entries are a subset of the non-zero entries in $x'$.

**Axioms and Theorem 1.** SHAP values are the unique solution that satisfies three desirable axioms:

- **Local accuracy (additivity):** The explanation model output equals the original model prediction:
- **Missingness:** If a feature is missing (i.e., $x_i' = 0$), then its attribution is zero: $\phi_i = 0$.
- **Consistency:** If a model changes such that a feature's contribution increases (or stays the same) for all subsets, then its SHAP value should not decrease.

Theorem 1 in [2] states that SHAP values are the only additive feature attribution method that satisfies all three of these properties. For a general audience, this means that SHAP assigns credit to each feature in a way that is provably fair, faithful to the model, and responsive to any structural improvements in how the model uses a feature. It guarantees that the sum of the parts (the individual feature attributions) always matches the whole (the model's prediction), and that features can't be given credit for outcomes they didn't influence.

**Kernel SHAP.** Kernel SHAP is a model-agnostic estimator that approximates SHAP values by fitting a weighted linear model over $z'$.

$$\pi_{x'}\left(z'\right) = \frac{(M-1)}{(M \text{ choose } |z'|)\,|z'|\,(M-|z'|)}$$

This kernel ensures that the solution aligns with the classical Shapley value formulation by weighting feature subsets appropriately in a least-squares regression framework. The loss function is given by:

$$\mathcal{L}\left(f, g, \pi_{x'}\right) = \sum_{z' \in Z} \left[f\left(h_x\left(z'\right)\right) - g\left(z'\right)\right]^2 \pi_{x'}\left(z'\right)$$

which penalizes discrepancies between the model's true output and the linear explanation model while incorporating the derived kernel weights. This kernel weighting ensures that the regression satisfies the three SHAP axioms. Kernel SHAP is based on LIME [4] but derives the loss, regularization, and weighting analytically rather than heuristically.

**TreeSHAP.** TreeSHAP is a specialized algorithm that computes SHAP values exactly in polynomial time for decision tree models (e.g., XGBoost, LightGBM). Instead of sampling subsets, it recursively traverses each tree and computes the expected contribution of each feature conditioned on its appearance along the tree paths [5].

The algorithm computes:

$$\phi_i = \mathbb{E}_{z' \sim \text{coalitions without } i}\left[f_x(z' \cup \{i\}) - f_x(z')\right]$$

TreeSHAP calculates these expectations analytically by tracking feature dependencies at each split node and propagating path weights and outputs through the tree structure. It avoids sampling and respects feature dependencies implicitly encoded in the tree, making it accurate and scalable for high-dimensional tabular data.

**DeepSHAP.** DeepSHAP extends SHAP to deep neural networks by combining DeepLIFT's backpropagation rules with SHAP's theoretical foundation [5, 6]. It approximates SHAP values by decomposing the output of the network $f(x)$ relative to a reference input $x'$:

$$\phi_i \approx (x_i - x'_i) \cdot \frac{\partial f}{\partial x_i}\Big|_{x'}$$

DeepSHAP assigns attributions layer by layer by computing multipliers for each neuron, then propagating these multipliers backwards from the output to the input. These multipliers satisfy the "summation-to-delta" property:

$$\sum_{i=1}^{M} \phi_i = f(x) - f(x')$$

While DeepSHAP is only an approximation to the true SHAP values (because it relies on linearity assumptions between layers and independent inputs), it is much faster than Kernel SHAP and allows interpretation of deep models like MLPs and CNNs. In our study, we use KernelSHAP and DeepSHAP to interpret multilayer perceptron models trained to predict RNA Polymerase II occupancy. In parallel, we compare these results with TreeSHAP outputs derived from gradient-boosted tree models, leveraging their different assumptions about feature independence and internal structure. This multi-model approach improves the robustness of SHAP-based attributions and helps us identify biologically meaningful regulatory features with greater confidence.

#### Sequencing data preprocessing

As described in Materials and Methods: ChIP-seq data analysis, the raw sequencing data for ChIP-seq experiments were downloaded from the Gene Expression Omnibus (GEO) and processed using a custom snakemake workflow (Supplementary Methods Fig 1A). The workflow includes the following steps:

1. Downloading raw sequencing data from GEO.
2. Aligning reads to the reference genome using Bowtie2 [7].
3. Removing reads mapping to ENCODE blacklisted regions to improve data quality.
4. Calling peaks and generating signal tracks using MACS2 [8].
5. Generating treatment and control pileup tracks.
6. Calculating fold enrichment tracks using the `bdgcmp` command.
7. Quantifying the signal using bigWigAverageOverBed [9] to calculate the mean signal across defined genomic regions, such as promoters and gene bodies.

For the Materials and Methods: RNA-seq and TT-seq data analysis, the raw sequencing data for RNA-seq and TT-seq experiments were also downloaded from GEO and processed using a custom snakemake workflow (Supplementary Methods Fig 1B). The workflow includes the following steps:

1. Downloading raw sequencing data from GEO.
2. Aligning reads to the reference genome using Bowtie2 [7].
3. Identifying and removing PCR duplicates using SAMtools markup [10] to eliminate artifacts due to library preparation.
4. Quantifying gene-level read counts using featureCounts [11] from the Subread package.
5. Performing differential gene expression analysis using DESeq2 [12] in R, applying default parameters to identify significantly differentially expressed genes.

The DAGs (Directed Acyclic Graphs) of the snakemake workflows are shown in Supplementary Methods Fig 1. The complete workflow is publicly available on GitHub at https://github.com/kashyapchhatbar/SHAP-analysis.

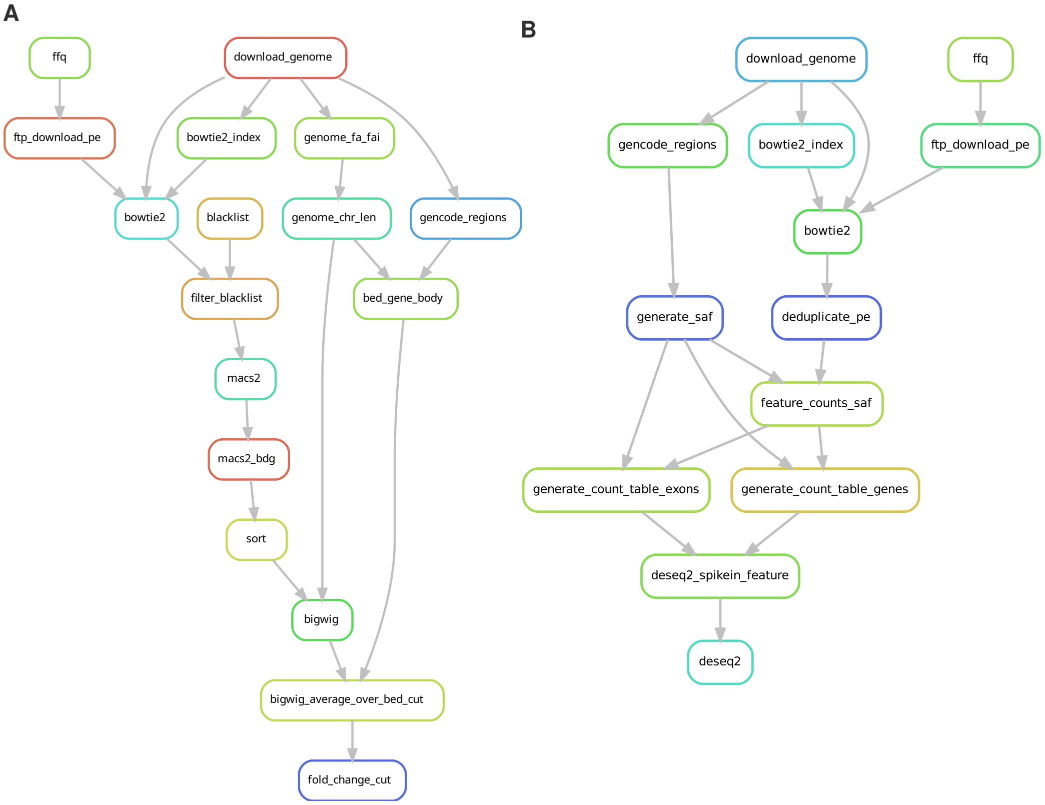


**Supplementary Methods Fig 1.** DAG of the snakemake workflow for ChIP-seq data analysis.

#### Hyperparameter optimisation

The hyperparameter optimisation for the MLPRegressor and XGBoostRegressor models was performed using the Optuna library [13] as described in Materials and Methods: Data preparation and modelling. The following tables summarise the hyperparameters of the models trained on Hughes et al. [14], Dobrinić et al. [15] and Wang et al. [16] data.

**Supplementary Methods Table 1.** Hyperparameters of the Scikit-learn MLPRegressor model trained on Hughes et al. [14] data.

| l2_regularisation | hidden_layer_sizes_1 | hidden_layer_sizes_2 | activation |
|---|---|---|---|
| 7.5167e-03 | 64 | 57 | relu |
| 1.3269e-05 | 62 | 56 | relu |
| 1.3797e-04 | 54 | 58 | relu |
| 9.6820e-05 | 45 | 61 | relu |
| 2.4365e-05 | 64 | 61 | relu |

**Supplementary Methods Table 2.** Hyperparameters of the PyTorch MLP model trained on Hughes et al. [14] data.

| learning_rate_init | weight_decay | hidden_layer_sizes_1 | hidden_layer_sizes_2 |
|---|---|---|---|
| 2.6827e-03 | 2.6086e-04 | 53 | 37 |
| 1.7020e-03 | 2.5392e-04 | 58 | 59 |
| 2.6403e-03 | 1.0215e-04 | 48 | 40 |
| 8.1475e-04 | 1.0007e-04 | 61 | 61 |
| 3.8772e-03 | 2.3777e-04 | 31 | 37 |

**Supplementary Methods Table 3.** Hyperparameters of the XGBoostRegressor model trained on Hughes et al. [14] data.

| learning_rate_init | tree_method | max_depth |
|---|---|---|
| 9.1785e-02 | exact | 6 |
| 7.9558e-02 | exact | 6 |
| 5.4488e-02 | exact | 6 |
| 6.5311e-02 | exact | 6 |
| 7.4898e-02 | exact | 7 |

**Supplementary Methods Table 4.** Hyperparameters of the Scikit-learn MLPRegressor model trained on Dobrinić et al. [15] data.

| l2_regularisation | hidden_layer_sizes_1 | hidden_layer_sizes_2 | activation |
|---|---|---|---|
| 4.5145e-05 | 61 | 64 | relu |
| 1.2066e-03 | 56 | 55 | relu |
| 1.4178e-04 | 62 | 64 | relu |
| 3.8390e-04 | 47 | 63 | relu |
| 1.2337e-03 | 60 | 58 | relu |

**Supplementary Methods Table 5.** Hyperparameters of the PyTorch MLP model trained on Dobrinić et al. [15] data.

| learning_rate_init | weight_decay | hidden_layer_sizes_1 | hidden_layer_sizes_2 |
|---|---|---|---|
| 6.9995e-04 | 2.1842e-04 | 61 | 51 |
| 2.5050e-03 | 4.3886e-04 | 56 | 23 |
| 2.2141e-03 | 1.4781e-04 | 60 | 27 |
| 4.0268e-03 | 2.5540e-04 | 46 | 62 |
| 6.9362e-03 | 3.9068e-04 | 42 | 51 |

**Supplementary Methods Table 6.** Hyperparameters of the XGBoostRegressor model trained on Dobrinić et al. [15] data.

| learning_rate_init | tree_method | max_depth |
|---|---|---|
| 7.4072e-02 | hist | 7 |
| 9.2142e-02 | hist | 6 |
| 6.2993e-02 | hist | 8 |
| 6.6056e-02 | hist | 7 |
| 6.4559e-02 | hist | 8 |

**Supplementary Methods Table 7.** Hyperparameters of the Scikit-learn MLPRegressor model trained on Wang et al. [16] data.

| l2_regularisation | hidden_layer_sizes_1 | hidden_layer_sizes_2 | activation |
|---|---|---|---|
| 1.9380e-02 | 60 | 61 | relu |
| 3.7036e-02 | 57 | 32 | relu |
| 2.7613e-02 | 48 | 59 | relu |
| 2.7805e-02 | 57 | 59 | relu |
| 1.6344e-02 | 63 | 56 | relu |

**Supplementary Methods Table 8.** Hyperparameters of the PyTorch MLP model trained on Wang et al. [16] data.

| learning_rate_init | weight_decay | hidden_layer_sizes_1 | hidden_layer_sizes_2 |
|---|---|---|---|
| 2.0964e-03 | 2.7503e-04 | 64 | 20 |
| 5.0577e-04 | 4.7558e-04 | 56 | 33 |
| 4.0241e-03 | 2.2214e-04 | 52 | 63 |
| 1.5923e-03 | 1.4617e-04 | 59 | 29 |
| 3.9265e-03 | 4.4898e-04 | 62 | 27 |

**Supplementary Methods Table 9.** Hyperparameters of the XGBoostRegressor model trained on Wang et al. [16] data.

| learning_rate_init | tree_method | max_depth |
|---|---|---|
| 5.0245e-02 | exact | 8 |
| 6.5811e-02 | hist | 7 |
| 6.7234e-02 | exact | 7 |
| 6.7051e-02 | hist | 7 |
| 5.7773e-02 | exact | 9 |

#### Model training evaluation

The training process was performed again involving fitting the models to the training data and evaluating their performance using the best hyperparameters found during the optimisation. Supplementary Methods Fig 2 illustrates the training and validation MSE (mean squared error) losses and $R^2$ scores at each epoch for the PyTorch MLP models.

**A** Metrics for PyTorch MLP model trained on Hughes et al. data set

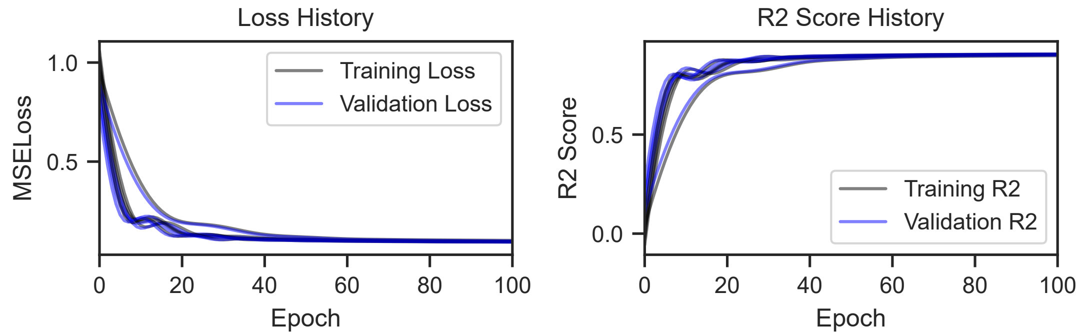


**B** Metrics for PyTorch MLP model trained on Dobrinić et al. data set

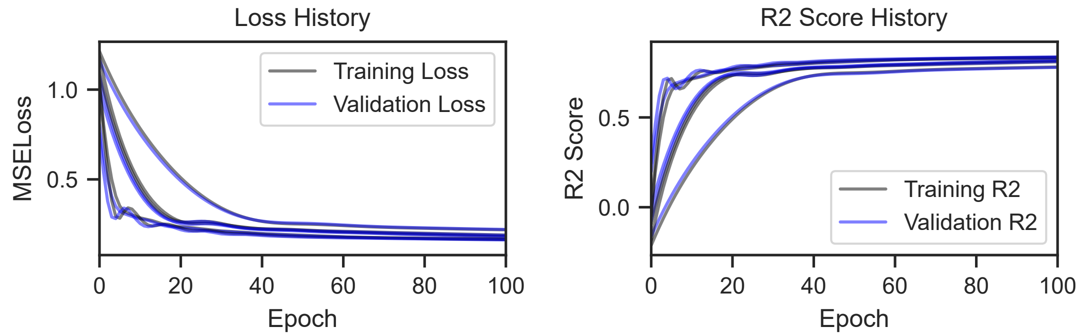


**C** Metrics for PyTorch MLP model trained on Wang et al. data set

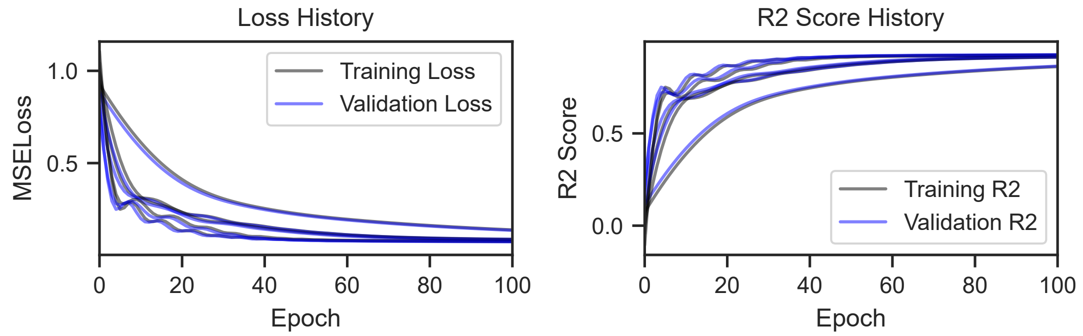


**Supplementary Methods Fig 2.** Training and validation MSE losses and $R^2$ scores for the PyTorch MLP models trained on A) Hughes et al. [14] B) Dobrinić et al. [15] C) Wang et al. [16] data.

#### Feature correlation

To gauge how much shared signal exists among our chromatin features and whether this threatens interpretability, we calculated pair-wise Pearson correlations for every dataset used in modelling. The resulting matrices were rendered as clustered heatmaps to highlight groups of features that move together. For Hughes et al. [14] minimal three-feature model (SET1A, ZC3H4, H3K4me3), the results shows substantial positive correlations. Promoter (p) signals cluster separately than gene body (gb) signals for SET1A and ZC3H4, supporting our decision to model the two regions separately (Supplementary Methods Fig 3A). In Dobrinic et al. [15], promoter and gene body signals form separate clusters with an isolated H3K27me3 gene body signal (Supplementary Methods Fig 3B). We see high multi-colinearity in Wang et al. [16] data and separate clusters of promoter and gene body (Supplementary Methods Fig 3C). Mean absolute SHAP values across all splits are highly consistent, and the rank order of feature importance was identical across DeepSHAP, KernelSHAP and TreeSHAP. Because TreeSHAP leverages the tree structure, it is immune to the independence assumption that underlies KernelSHAP, providing an orthogonal check on the explanations. Together, these tests show that SHAP still identifies biologically meaningful drivers even in the presence of moderate to high feature correlations.

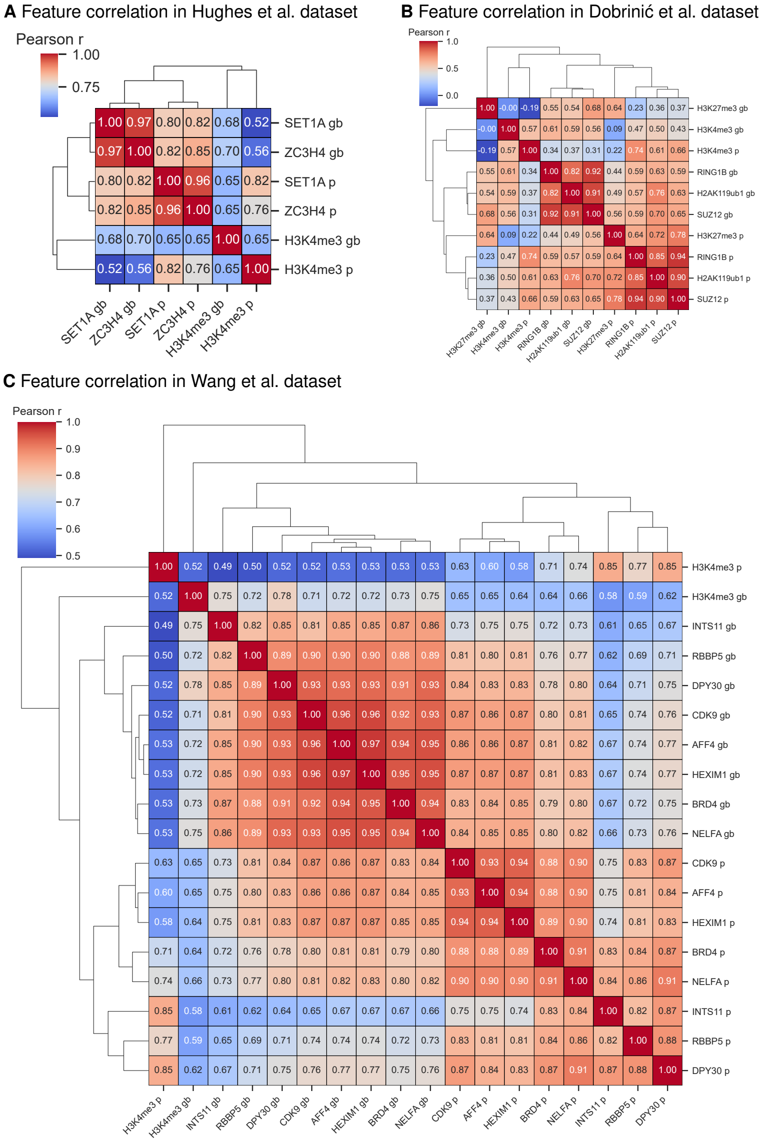


**Supplementary Methods Fig 3.** Correlation structure of chromatin features across A) Hughes et al. [14] B) Dobrinić et al. [15] and C) Wang et al. [16] data. The input data frames of training sets were used to calculate the pearson correlation coefficient matrices for all three data sets and is visualised in clustered heatmaps. Promoter=p, Gene body=gb.

#### References


1. Linardatos P, Papastefanopoulos V, Kotsiantis S. Explainable AI: A Review of Machine Learning Interpretability Methods;23(1):18. doi:10.3390/e23010018.

2. Lundberg SM, Lee SI. A unified approach to interpreting model predictions. In: Advances in neural information processing systems. vol. 30. Curran Associates, Inc.;.Available from: https://papers.nips.cc/paper_files/paper/2017/hash/8a20a8621978632d76c43dfd28b67767-Abstract.html.

3. Shapley LS. 17. A Value for n-Person Games. In: Kuhn HW, Tucker AW, editors. Contributions to the Theory of Games (AM-28), Volume II. Princeton University Press;. p. 307–318. Available from: https://www.degruyter.com/document/doi/10.1515/9781400881970-018/html.

4. Ribeiro MT, Singh S, Guestrin C. "Why should I trust you?": Explaining the predictions of any classifier. In: Proceedings of the 22nd ACM SIGKDD international conference on knowledge discovery and data mining. KDD '16. Association for Computing Machinery;. p. 1135–1144. Available from: https://dl.acm.org/doi/10.1145/2939672.2939778.

5. Lundberg SM, Erion GG, Lee SI. Consistent Individualized Feature Attribution for Tree Ensembles;. Available from: http://arxiv.org/abs/1802.03888.

6. Shrikumar A, Greenside P, Kundaje A. Learning Important Features Through Propagating Activation Differences;. Available from: http://arxiv.org/abs/1704.02685.

7. Langmead B, Salzberg SL. Fast gapped-read alignment with Bowtie 2;9(4):357–359. doi:10.1038/nmeth.1923.

8. Zhang Y, Liu T, Meyer CA, Eeckhoute J, Johnson DS, Bernstein BE, et al. Model-based analysis of ChIP-seq (MACS);9(9):R137. doi:10.1186/gb-2008-9-9-r137.

9. Kent WJ, Zweig AS, Barber G, Hinrichs AS, Karolchik D. BigWig and BigBed: enabling browsing of large distributed datasets;26(17):2204–2207. doi:10.1093/bioinformatics/btq351.

10. Li H, Handsaker B, Wysoker A, Fennell T, Ruan J, Homer N, et al. The sequence alignment/map format and SAMtools;25(16):2078–2079. doi:10.1093/bioinformatics/btp352.

11. Liao Y, Smyth GK, Shi W. featureCounts: an efficient general purpose program for assigning sequence reads to genomic features;30(7):923–930. doi:10.1093/bioinformatics/btt656.

12. Love MI, Huber W, Anders S. Moderated estimation of fold change and dispersion for RNA-seq data with DESeq2;15(12):550. doi:10.1186/s13059-014-0550-8.

13. Akiba T, Sano S, Yanase T, Ohta T, Koyama M. Optuna: A Next-generation Hyperparameter Optimization Framework;. Available from: http://arxiv.org/abs/1907.10902.

14. Hughes AL, Szczurek AT, Kelley JR, Lastuvkova A, Turberfield AH, Dimitrova E, et al. A CpG island-encoded mechanism protects genes from premature transcription termination;14(1):726. doi:10.1038/s41467-023-36236-2.

15. Dobrinić P, Szczurek AT, Klose RJ. PRC1 drives Polycomb-mediated gene repression by controlling transcription initiation and burst frequency;28(10):811–824. doi:10.1038/s41594-021-00661-y.

16. Wang H, Fan Z, Shliaha PV, Miele M, Hendrickson RC, Jiang X, et al. H3K4me3 regulates RNA polymerase II promoter-proximal pause-release;615(7951):339–348. doi:10.1038/s41586-023-05780-8.
